# A Versatile Vector System for the Fast Generation of Knock-in Cell Lines with CRISPR

**DOI:** 10.1101/2020.02.06.927384

**Authors:** Oscar Perez-Leal, Jonathon Nixon-Abell, Carlos A. Barrero, John Gordon, Mario C. Rico

## Abstract

Until recent advancements in genome editing via CRISPR/Cas9 technology, understanding protein function typically involved artificially overexpressing proteins of interest. Despite that CRISPR/Cas9 has ushered in a new era of possibilities for modifying endogenous genes with labeling tags (knock-in) to more accurately study proteins under physiological conditions, the technique is largely underutilized due to its tedious, multi-step process. Here we outline a homologous recombination system (FAST-HDR) to be used in combination with CRISPR/Cas9 that significantly simplifies and accelerates this process while introducing multiplexing to allow live-cell studies of 3 endogenous proteins within the same cell line. Furthermore, the recombination vectors are assembled in a single reaction that is enhanced for eliminating false positives and reduces the overall creation time for the knockin cell line from ^~^8 weeks to <15 days. Finally, the system utilizes a modular construction to allow for seamlessly swapping labeling tags to ensure flexibility according to the area under study. We validated this new methodology by developing advanced cell lines with 3 fluorescent-labeled endogenous proteins that support high-content phenotypic drug screening without using antibodies or exogenous staining. Therefore, Fast-HDR cell lines provide a robust alternative for studying multiple proteins of interest in live cells without artificially overexpressing labeled proteins.

## INTRODUCTION

The recent discovery and development of CRISPR/Cas9 endonucleases for targeting specific DNA sequences in living cells have advanced genetic engineering in eukaryotic organisms^1^. While the more common application is to conduct gene alteration for loss-of-function studies (gene knockouts), modifying DNA by insertions of new genetic information (gene knock-ins) is particularly useful for tagging endogenous genes, enabling the study of biochemical and dynamic properties of encoded proteins. To create cell lines with tagged endogenous genes, a donor DNA template containing the genetic information must first be created, then inserted – typically using homology-directed repair (HDR)^2^ – after cutting the DNA via double-strand break (DSB) with CRISPR/Cas9 endonucleases. The donor templates used to promote HDR are large plasmids containing homologous recombination arms for the 5’ and 3’ ends of the cut site^3^. These donor plasmids also contain the genetic sequences of the inserted protein markers as well as recombinant cassettes for the expression of a resistance gene to facilitate the isolation of modified cell clones. Currently, access to plasmid backbones to develop HDR donor templates is limited to a few commercial plasmids or a small selection of plasmids available in a nonprofit plasmid repository^4,5^. These plasmids typically only allow tagging with one specific protein marker and also limit the selection of modified cell lines to only one type of eukaryotic resistance gene^4,5^. Therefore, most researchers interested in knock-in tagging projects follow guidelines to build their own donor template plasmids^6^, which is laborious and requires a multistep process that can take more than three weeks to complete^7^.

We proposed that redesigning the donor template plasmid to facilitate the construction of advanced homologous recombination plasmids could circumvent these limitations while simplifying and accelerating the overall cell-line creation process if the vector system included the following characteristics: 1) The backbone system should allow the insertion of any protein tag into a gene of interest, 2) it should allow the insertion of tags in at least 3 targets in the same cell line (multiplexing) to facilitate the use of high-content imaging analysis, 3) it should allow the insertion of corresponding resistance genes to enable pure cell selection for the protein tag of choice, 4) it should ideally, construct the above in a single-step reaction that significantly shortens the overall creation time of the donor template.

Here, we report the development of a system that combines a novel set of advanced donor plasmids for HDR and a streamlined workflow that simplifies and enhances the process for developing next generation cell lines with tagged endogenous genes. Furthermore, the construction of these plasmids are achieved in a single-step reaction that can be completed in only 1 day versus ^~^21 days with current methodologies^7^, while also achieving additional benefits downstream that further accelerate the overall cell-line creation process to <15 days compared to ^~^8 weeks using current methodologies^7^. In addition to speed, the system also enhances the complexity of the cell models that can be generated by supporting homozygous tagging and introducing multiplexing of up to 3 targets. Thus, our method requires neither staining nor fixing to perform visual analysis, as demonstrated using high-content phenotypic drug screening. This represents a breakthrough in live-cell studies for understanding cells’ biological functions by simplifying, accelerating and enhancing the process for utilizing CRISPR knock-ins for tagging endogenous genes.

## RESULTS

### Construction of a plasmid backbone system for the fast assembly of donor templates

The generation of homologous recombination donor templates for the knock-in of large labeling tags requires the assembly of a recombination cassette containing four main elements. 1) A 5’ target-specific recombination arm sequence, 2) the sequence of a labeling tag, 3) an antibiotic resistance gene for selection of modified cells and 4) a 3’ target-specific recombination arm sequence. Traditionally, these elements are selected and assembled in a multi-step process that can take up to 3 weeks to complete for every target gene^7^. In order to overcome these limitations, we designed the FAST-HDR system (Fig. 1) as a series of backbone plasmids that facilitates the creation of donor templates for inserting multiple types of large labeling tags. Each backbone plasmid with a specific labeling tag allows the incorporation of both the 5’ and 3’ recombination arms in a single enzymatic reaction, thus avoiding the traditional multi step assembly process.

**Figure 1.**
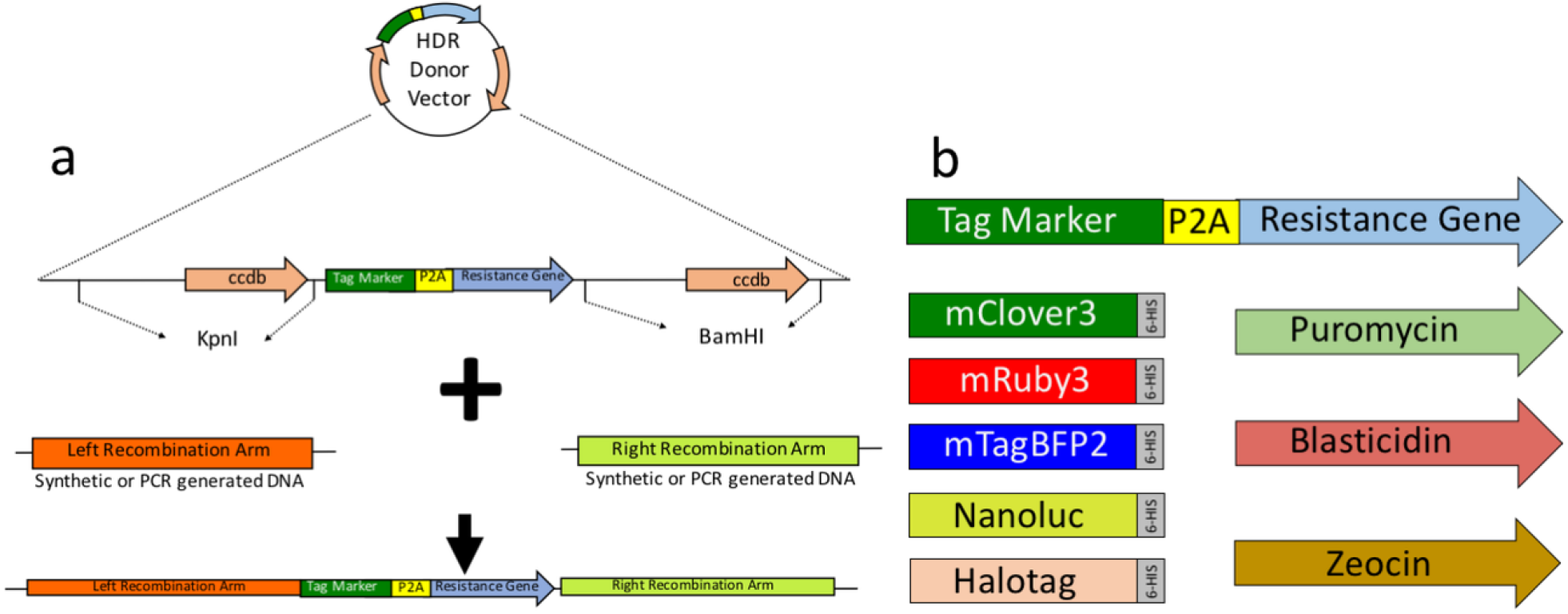
The FAST-HDR vector system. (a) Diagram of the general components of the plasmid that facilitates the rapid construction of donor templates for homologous recombination. The plasmid is first digested with restriction enzymes to cut the segments with CddB cassettes (in this case, KpnI and BamHI). Previously designed synthetic recombination arms are cloned into the digested backbone plasmid using the Gibson assembly method. (b) Diagram of the protein tags and selection antibiotics that are included in the FAST-HDR system.

To construct the FAST-HDR system, we combined three existing technologies for accelerating gene cloning. First, double cloning of the 5’ and 3’ recombination arms was performed in a single step by promoting the replacement of two copies of a lethal gene for *E.coli, ccdB*^8^, thus allowing only the growth of colonies with the correct insertions of the arms. Second, the 5’ and 3’ recombination arms were provided as sequence-verified synthetic dsDNA. Third, the Gibson assembly method^9^ facilitated the correct insertion of the 5’ and 3’ recombination arms in a single reaction (Fig. 1A). Thus, this streamlined method reduces the time for generating an HDR plasmid from ^~^3 weeks to only 1 day and requires less than 30 minutes of hands-on time. In addition, the system is now set up for allowing the insertion of multiple protein tags for downstream applications, such as fluorescence microscopy, luminescence, and protein purification (Fig. 1B). The current system is preconfigured with 5 different protein tags, and 3 eukaryotic antibiotics. The system can also be easily modified to include any protein tag or eukaryotic resistance gene that a researcher might need.

The selection of pure cell clones after gene editing typically requires the use of equipment for cell sorting^3^, which due to its high expense, is not available in every laboratory. Alternatively, the isolation of modified clones can be performed by cell selection with resistance cassettes against eukaryotic antibiotics, followed by single cell dilution – a process that with existing methods can take several weeks^3,7^. One additional benefit of the FAST-HDR vector system is that these plasmids allow the in-frame expression of any eukaryotic antibiotic resistance gene for rapid selection of only modified cells.

### Fast generation of cell lines with knock-in labeling tags

We tested the FAST-HDR plasmids by generating multiple cell lines that highlight the benefits of this system and present a broad spectrum of applications. First, we created 3 plasmid backbones to allow the tagging of endogenous genes with red fluorescent protein mRuby3^10^, the green fluorescent protein mClover3^10^, and the blue fluorescent protein mTagBFP2^11^. Each backbone plasmid was also built to contain a different eukaryotic resistance gene (zeocin, puromycin, and blasticidin respectively). Next, we targeted HEK293T cells with CRISPR-Cas9 and the HDR plasmids to modify endogenous genes encoding proteins with well-defined subcellular localizations. In particular, we C-terminally tagged the genes encoding Histone 3.3 (H3.3, nucleus), β tubulin (microtubules), and ATP synthase subunit β (ATP5B; mitochondria), with mRuby3, mClover3, and mTagBFP2, respectively (Fig. 2A-C and Supplementary Fig. 1). In all cases, the new HDR plasmids and workflow (Supplementary Fig. 2) allowed the selection of modified cell lines between 9 and 12 days after assembling the HDR plasmids.

**Figure 2.**
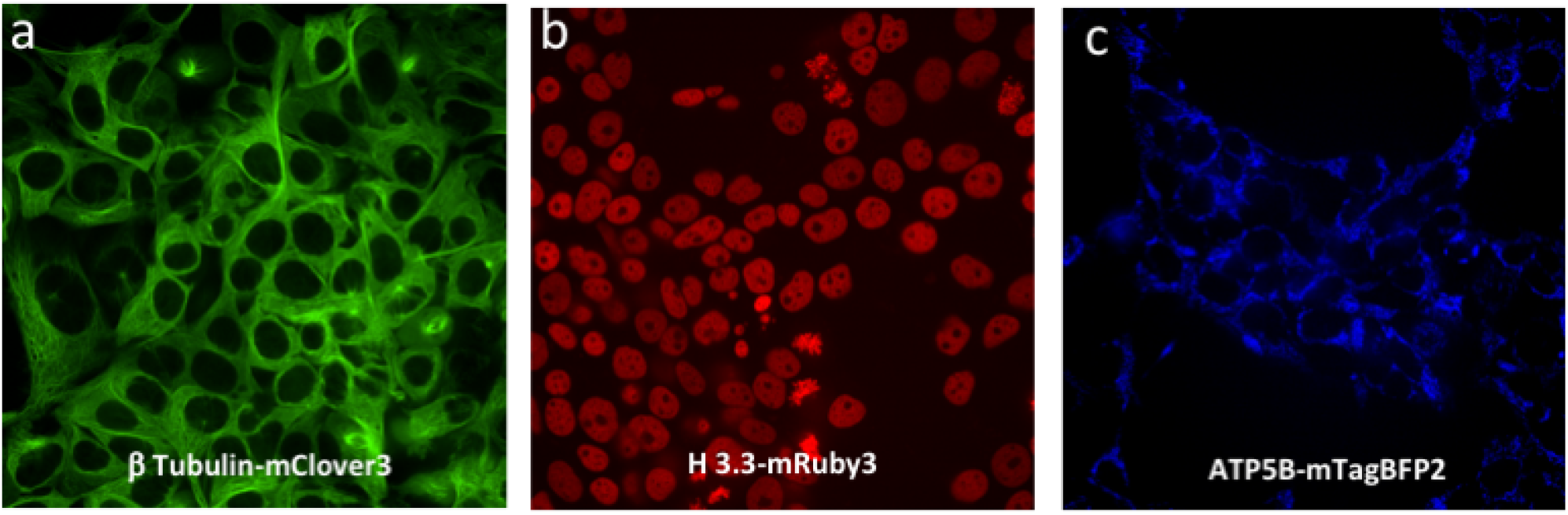
Generation of Cell lines with knock-in fluorescent tags with the FAST-HDR system. (a-c) HEK293T cell lines with labeling of (a) beta tubulin with mClover3, (b)H3.3 with mRuby3, and (c) ATP5B with mtagBFP2.

### The FAST-HDR vector system facilitates generating homozygous tagging of target genes

To test whether the FAST-HDR system can be used to generate homozygous labeled cell lines, we cloned the recombination arms into two backbone plasmids with different resistance genes for dual antibiotic selection. We targeted near-diploid HTC116 cells^12^ with a combination of CRISPR-Cas9 and two HDR donor plasmids to tag the transcriptional regulator of the antioxidant response, NRF2 (nuclear factor (erythroid-derived 2)-like 2), with NanoLuc luciferase^13^ at the C-terminus. NRF2 is a low-abundance protein that is tightly regulated by oxidative stress^14^. We confirmed the tagging of both alleles of *NRF2* (Supplementary Fig. 1D). This cell line facilitated detection of NRF2 levels in real time in response to an inducer of ox-idative stress, tert-butyl hydroperoxide (TBHP)^15^, or to two compounds that prevent the rapid degradation of NRF2, sulforaphane (SFN)^16^ and tert-butyl hydroquinone (TBHQ)^17^ (Fig. 3). Therefore, the FAST-HDR system can be used to homozygously tag proteins of interest to study their behavior under different conditions.

**Figure 3.**
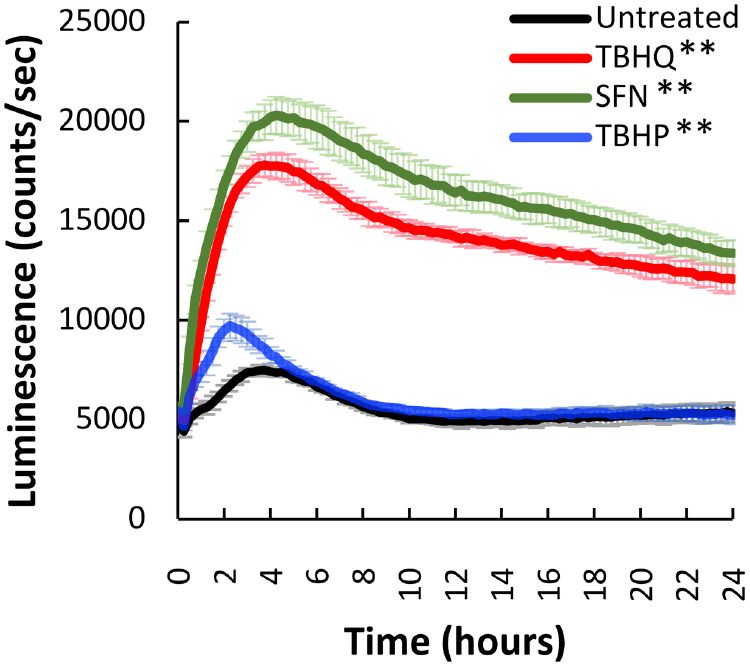
Generation of a cell line with knock-in endogenous labeling of Nrf2 with Nanoluc luciferase. Time-course luminescence produced by an HCT116 cell line with homozygous tagging of the NRF2 protein with NanoLuc, in the presence or absence of terbutylhydroquinone (TBHQ), sulforaphane (SFN), and terbutylhydroperoxide (TBHP). Data shown are the average and s.d of six independent experiments. Differences were analyzed by twoway ANOVA and P < 0.05 was considered statistically significant. ** P < 0.0001

### Multiplexing knock-in insertions of three labeling tags with the FAST-HDR vector system

An additional feature of the FAST-HDR system is that these novel plasmids allow for multiplexing up to 3 targets in the same cell line, representing a significant breakthrough in studying multiple endogenous proteins simultaneously within cells. Furthermore, because these cell lines already have endogenous proteins labeled with fluorescent tags, they do not require the use of immunofluorescence or transfection-based insertion of exogenous plasmids to generate images for high-content imaging analysis. Thus, the system eliminates the additional time, cost, and effort required to detect the proteins of interest used in existing processes and allowing evaluation of cellular changes at different time points without the requirement for fixation.

To demonstrate the applicability of multiplexing with the FAST-HDR system, we generated two HEK293T cell lines each expressing three proteins tagged with different fluorescent reporters. Such cell lines, with multiple fluorescent tags, are an excellent tool for superresolution microscopy (Fig. 4) or live-cell confocal microscopy (Fig. 5). For example, we evaluated the growth of these cell lines by time-lapse microscopy after treatment with staurosporine, a well-known inducer of apoptosis (Fig. 5A and Supplementary Video 1), or paclitaxel, a microtubule-stabilizing drug (Fig. 5B and Supplementary Video 2).

**Figure 4.**
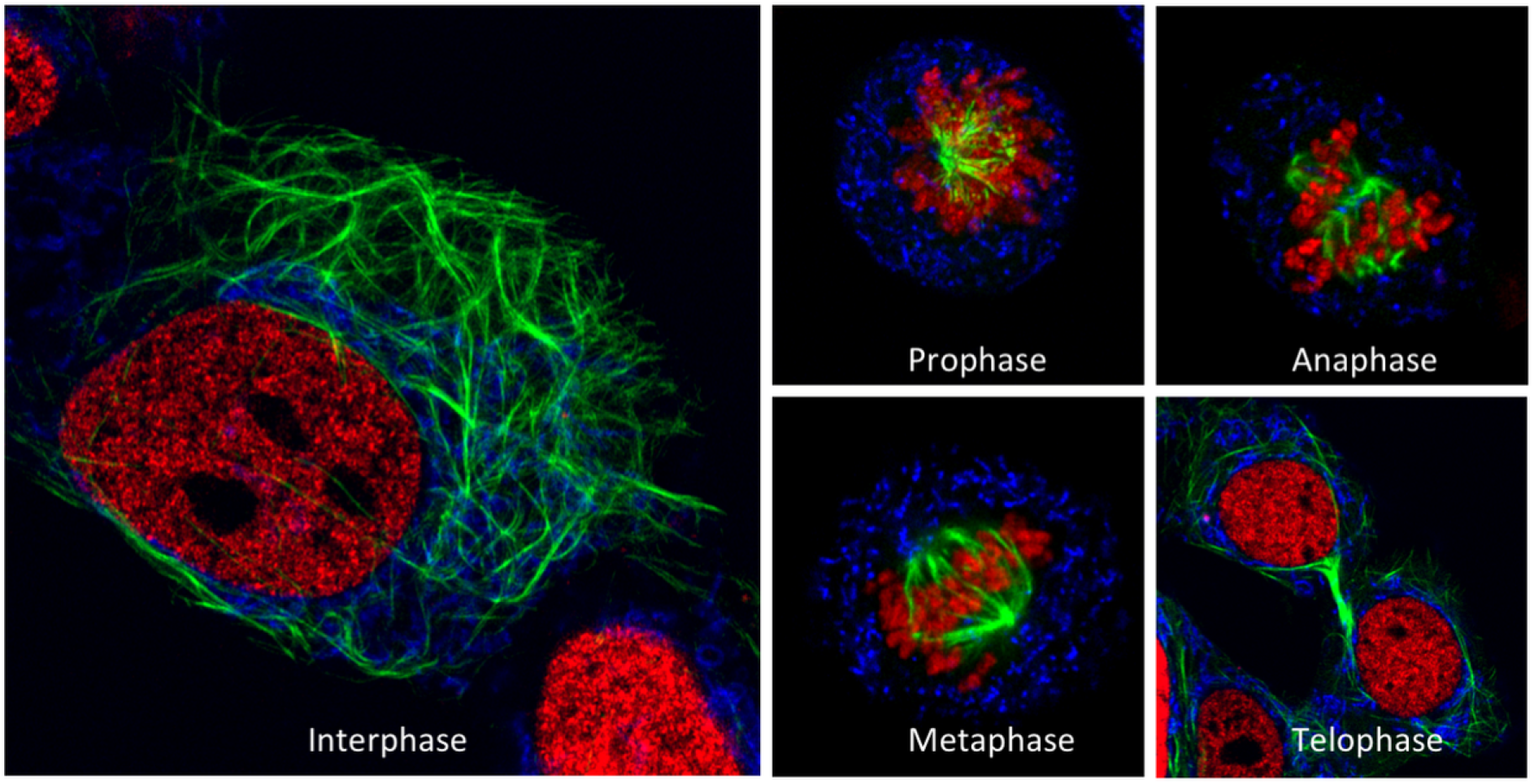
The FAST-HDR system facilitates the development of cell lines with multiple endogenously labeled genes. Evaluation of the mitotic states of HEK293T cells with nuclear (H3.3-mRuby3), microtubule (β tubulin-mClover3), and mitochondrial (ATP5B-mTagBFP2) labeling by superresolution confocal microscopy.

**Figure 5.**
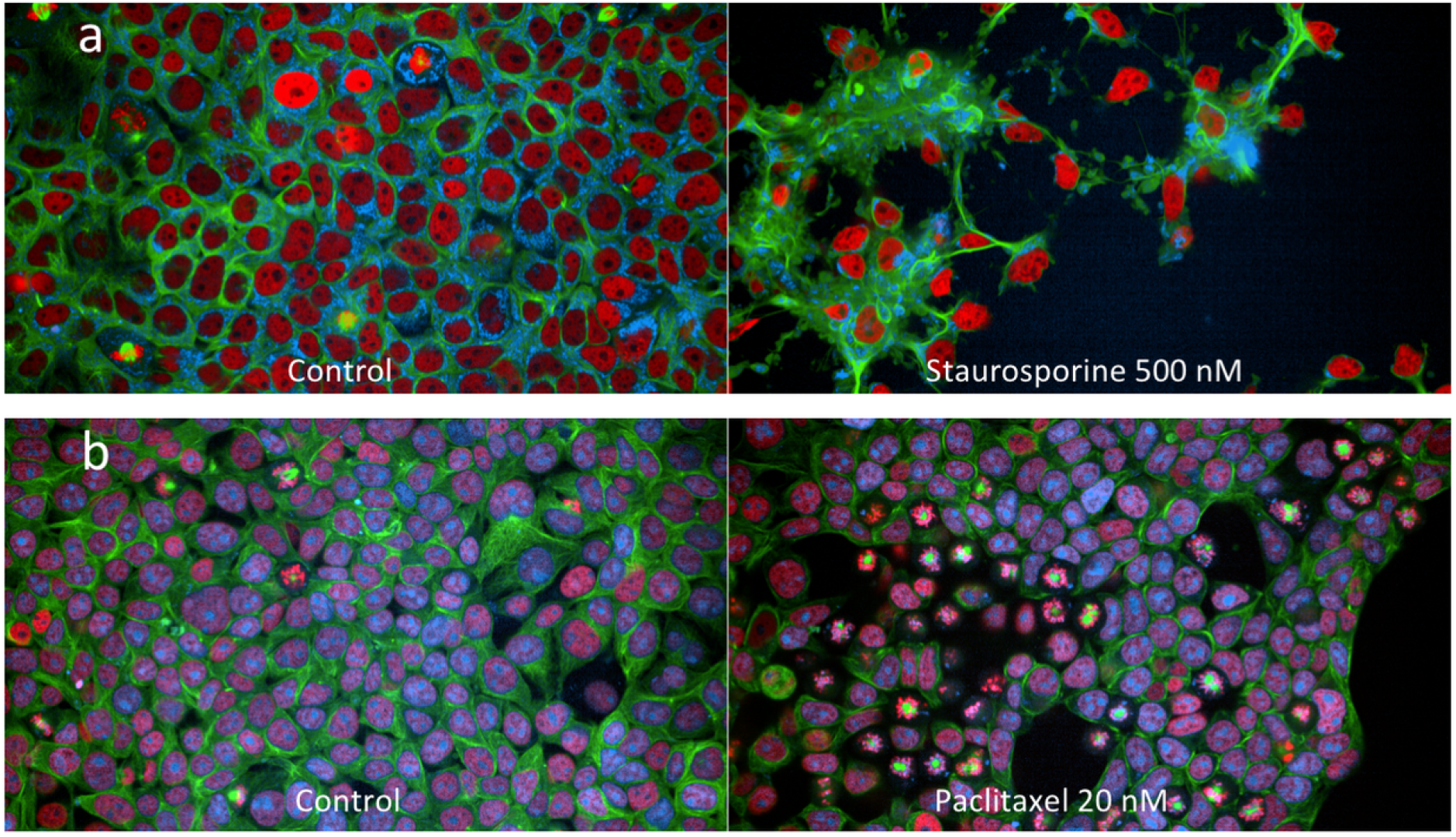
Use of multiplex cell lines for time-lapse confocal imaging without antibodies or chemical staining. (a) The growth of HEK293T cells with nuclear (H3.3-mRuby3), microtubule (β tubulin-mClover3), and mitochondrial (ATP5B-mTagBFP2) was evaluated by timelapse confocal microscopy for 13 h with and without an apoptosis inducer (staurosporine). The complete time-lapse sequence can be seen in Supplementary Video 1. (b) The growth of an HEK293T cell line with endogenous labeling of PARP1-mTagBFP2, H3.3-mRuby3, and β tubulin-mClover3 was evaluated by time-lapse confocal microscopy for 13 h with and without a microtubule-stabilizing drug (paclitaxel). The complete time-lapse sequence can be seen in Supplementary Video 2.

To further validate the applicability of these cell lines for high-content imaging drug screening, we created a HELA cell line for evaluating the accumulation of autophagic vesicles. For this, we tagged a nuclear protein (Histone 1-mtagBFP2), a cytoskeleton marker (β tubulin-mClover3), and a known autophagy receptor protein (SQSTM1-mRuby3) to screen for kinase inhibitors (with known targets) that induce the accumulation of cytoplasmic autophagic vesicles after 24 h of treatment (Supplementary Fig. 3). With this approach, we identified two kinase inhibitors (BI-6727 and CX-6258) that increase the number of autophagic vesicles similar to hydroxychloroquine, a known autophagy inhibitor (Supplementary Fig. 3B). We validated this finding by tracking the cytoplasmic autophagic vesicles accumulation of the same cell line with the PLK1 inhibitor (BI-6727) and the PIM kinases inhibitor (CX-6258) over 16 h (Fig. 6, Supplementary Video 3). Therefore, simultaneous tagging of up to three genes using the FAST-HDR system can be used to develop cellular models for high-content imaging analysis for drug discovery without the use of staining or immunofluorescence.

**Figure 6.**
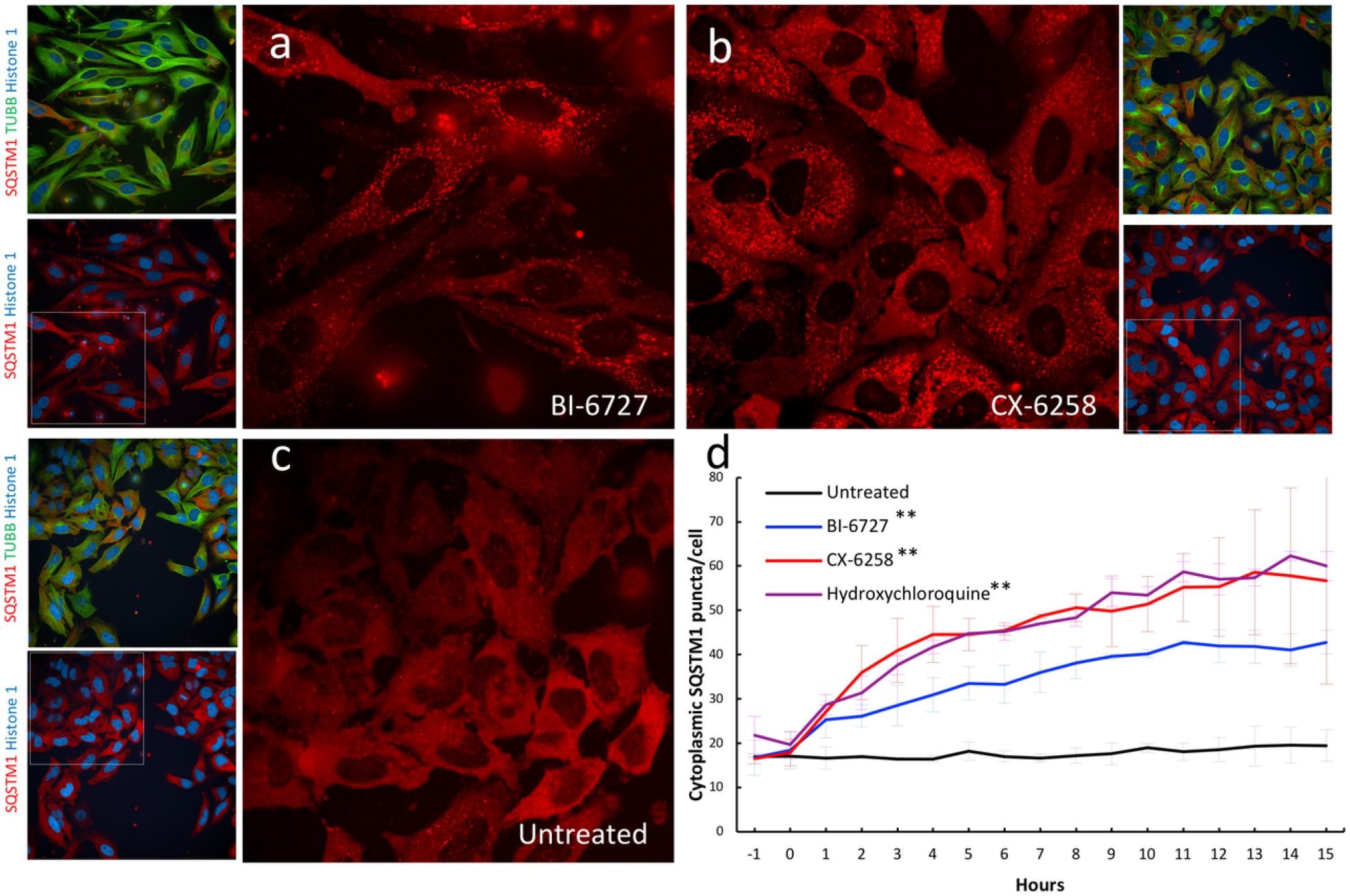
Validation of kinase inhibitors that induce the accumulation of autophagic vesicles in a triple labeled HELA cell line. A multiplex labeled HELA cell line with nuclear (Histone 1-m, TagBFP2), microtubule (Beta tubulin-mClover3), and the autophagic receptor protein p62 (GSTM1-mRuby3) was used to evaluate the cytoplasmic accumulation of autophagic vesicles after 16 hours of treatment with two kinase inhibitors (a) BI-6727 and (b) CX-6258 versus (c) Untreated cells. (d) The average number of autophagic vesicles per cell were counted every ten minutes and compared against untreated cells or cells treated with Hydroxychloroquine as a positive control. The complete time-lapse sequence can be seen in Supplementary Video 3. Data shown are the average and s.d. of three independent experiments. Differences were analyzed by two-way ANOVA and P < 0.05 was considered statistically significant. ** P < 0.0001

### The FAST-HDR vector system allows the use of any labeling tag for a target gene

Finally, the system allows for swappable tagging. That is, the single design of recombination arms can be inserted in any of the plasmids within the system to facilitate targeting the same gene with different tags. We confirmed that, by cloning one set of recombination arms into several HDR plasmids, the FAST-HDR system can be used to generate multiple cell lines with various modifications of the same target gene. We generated multiple HEK293T cell lines with the endogenous labeling of poly(ADP-ribose) polymerase 1 (PARP1) with three different fluorescent proteins (Fig. 7A).

**Figure 7.**
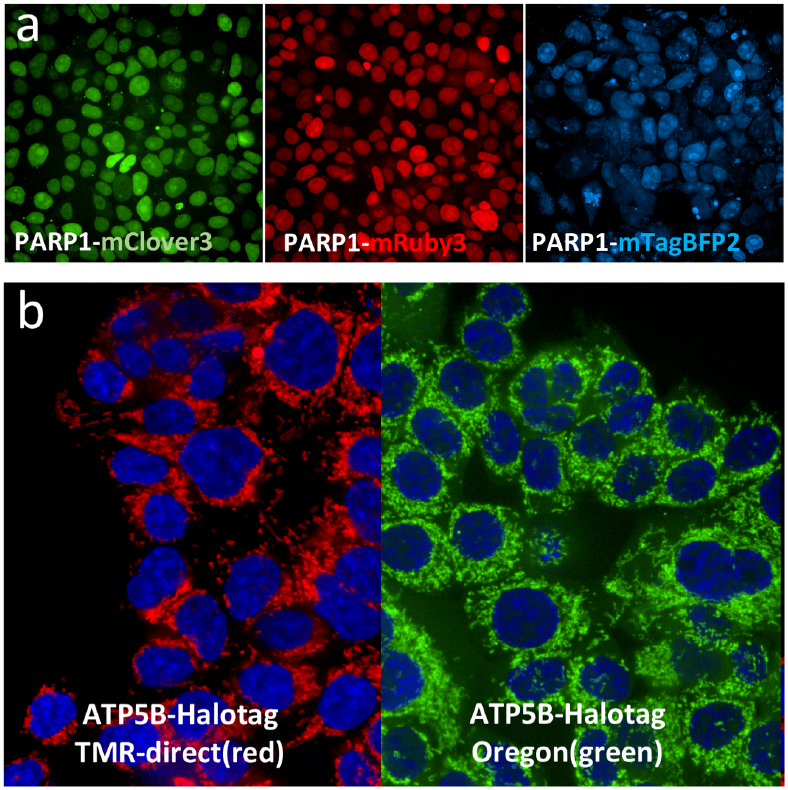
Using the FAST-HDR system to target an endogenous gene with multiple tags. (a) Three cell lines (HEK293T) with the PARP1 gene labeled with different fluorescent tags. (b) An HCT116 cell line with homozygous tagging of the ATP5B protein with the HaloTag. For protein visualization, the cells were treated with the HaloTag ligand TMR-direct (red) or Oregon Green and counterstained with Hoechst 33342 to detect the nucleus.

We also generated an HTC116 cell line with homozygous tagging of ATP5B with the purification/labeling tag HaloTag^18^ (Fig. 7B). For this purpose, we used the same recombination arm constructs as for the labeling of ATP5B with mTagBFP2 in Fig. 2C. Therefore, the FAST-HDR system offers the flexibility to label the same protein target with different tags, according to the application of the cell line.

## DISCUSSION

In this study, we have developed an efficient way to leverage CRISPR/Cas9 technology for genome editing, and specifically for inserting labeling tags into genes of interest (knockins) to track their proteins, as a robust alternative to traditional methods that typically involve artificially inducing recombinant protein overexpression. Though the technique is currently largely underutilized given the extensive preparation required to create a specific large donor template for each gene target under study^7^, the FAST-HDR vector system removes these critical barriers and provides a low-cost, single-step reaction for creating these donor templates that also enables more complex studies from the resulting cell lines.

The construction of the FAST-HDR system as a series of backbone plasmids that facilitate the creation of advanced donor templates offers several advantages over traditional HDR plasmids:

1. Each backbone plasmid with a specific labeling tag allows the incorporation of both the 5’ and 3’ recombination arms in a single enzymatic reaction, thus avoiding the traditional multi-step assembly process that discourages scientists from fully utilizing CRISPR technology for inserting large labeling tags – a feature that also increases the speed of obtaining a functional HDR donor template to only one day vs up to 3 weeks with other methodologies^7^.
2. The system allows the insertion of any protein tag into a gene of interest. The cassettes for the labeling tag and the mammalian selection marker were designed to be replaceable, thus allowing the easy generation of additional backbones for labeling genes with any type of protein tag and for selecting cell lines with multiple targeted genes (multiplexing). In the current version, we have included three fluorescent tags (mClover3, mRuby3, mtagBFP2), one luminescent tag (nanoluc) and one purification tag (halotag), as well as three mammalian selection antibiotics cassettes (puromycin, blasticidin, and zeocin). While the current FAST-HDR system requires the active expression of the target protein in the cell of interest to facilitate antibiotic selection, the generation of a plasmid template that includes a promoter cassette for the resistance gene would likely provide a method to overcome this in the future. Also, despite that the FAST-HDR system was tested with seven specific genes, there are currently no limitations that would prevent using the system for high throughput knock-in gene targeting. In fact, recent methodologies that facilitate high throughput knock-in gene targeting limit their use to a specific cell line and only allow targeting one gene with GFP^19^.
3. The FAST-HDR method allows the cloning of the genespecific recombination arms into any of the plasmid backbones in the system. This feature facilitates the development of donor templates for homozygous gene tagging via dual antibiotic selection. The same feature also allows the labeling of the same gene of interest with multiple tags and without increasing plasmid construction cost or time. This facilitates the study of genes of interest with complementary technologies; for example, it is possible to develop a cell line to study the gene of interest by fluorescent microscopy and also to develop an additional cell line to discover protein binding partners of the targeted protein by using covalent purification tags such as Halotag.

Altogether, this novel system accelerates the construction of cell lines for multiple applications, such as confocal and superresolution microscopy, the development of reporter cell lines for drug screening, and high-content imaging studies without the use of antibodies or exogenous fluorescent markers. The FAST-HDR system is therefore a more versatile methodology for developing endogenous tagging with CRISPR according to the needs of the researcher, as it works on any cell line, facilitates multiplexing and allows the use of any protein tag.

## MATERIALS AND METHODS

### FAST-HDR plasmid vector development

The HDR plasmids were built on a pUC57 backbone plasmid using synthetic DNA fragments (gBlocks from Integrated DNA Technologies, Skokie, Illinois, USA) and standard molecular biology techniques. The functional modules were organized from 5’ to 3’ as follows: first, a CcdB expression cassette flanked by KpnI recognition sites; second, a protein tag sequence flanked by EcoRI recognition sites and in frame with a P2A peptide sequence; third, a eukaryotic antibiotic resistance gene flanked by XbaI recognition sites; fourth, a MALAT 3’ sequence to provide mRNA stabilization^20^; and finally, a second CcdB expression cassette flanked by BamHI recognition sites. This design facilitates the rapid modification of the plasmid to include any protein tag or antibiotic resistance gene by digesting the plasmid with EcoRI or XbaI, respectively, and performing Gibson assembly with the desired DNA sequence.

### Cloning of recombination arms into the FAST-HDR system

The recombination arms were obtained as synthetic DNA fragments from Integrated DNA Technologies, and their sequences are provided in the Supplementary table 1. The recombination arms of each gene target were designed by selecting 400-660 nucleotides from the 5’ and 3’ sites around the expected DSB. The left recombination arm (5’ end) was designed to end with the last codon of the target gene to allow the in-frame expression of the protein tag and the antibiotic resistance gene. The recombination arms also included flanking sequences at the 5’ and 3’ ends to enable directional cloning by the Gibson assembly method, and all the recombination arms were compatible with all the plasmids described in this work. To allow insertion of the recombination arms into the FAST-HDR vector system, 500 ng of the plasmid was digested with KpnI and BamHI in a 20 μL reaction for 1 h using FastDigest enzymes (Thermo Fisher Scientific, Waltham, Massachusetts, USA). The recombination arms were resuspended in TE buffer at 25 ng/μL. Finally, the Gibson assembly reaction was performed for 1 h at 50°C using 2 μL of the digested plasmid, 1.5 μL of each recombination arm, and 5 μL of NEBuilder HiFi DNA Assembly Master Mix (New England Biolabs, Ipswich, Massachusetts, USA). Finally, 3 μL of the Gibson assembly reaction was used to transform chemically competent *E. coli* cells.

### CRISPR Cas9 and sgRNA guide expression

The plasmid expressing eSpCas9(1.1)^21^ was a gift from Feng Zhang (Addgene plasmid #71814). The U6 promoter was replaced with a tRNA promoter^22^ to facilitate the expression of sgRNA guides. For this purpose, the plasmid was digested with PscI and XbaI, and a synthetic DNA construct containing the tRNA sequence plus a CcdB selection cassette flanked by BbsI sites was inserted by Gibson assembly. This insertion facilitated fast cloning by Gibson assembly of the sequence encoding the sgRNA guides using 80 bp single-stranded DNA oligos. The 20 bp sgRNA guides for each target were selected with the CRISPR module of the Benchling online platform and are included in the Supplementary table 1.

### Cell culture

HEK293T, HCT116 and HELA cells were acquired from ATCC (ATCC, Manassas, Virginia, USA, Catalog numbers CRL-3216 and CCL-247 respectively). HEK293T and HELA cells were grown in Dulbecco’s modified Eagle’s medium (DMEM) supplemented with 10% fetal bovine serum (FBS) 100 U/ml penicillin, and 100 μg/ml streptomycin at 37°C and 5% CO_2_. HCT116 cells were grown in McCoy medium supplemented with 10% FBS and 100 U/ml penicillin, and 100 μg/ml streptomycin at 37°C and 5% CO_2_.

### Cell transfection and antibiotic selection

HEK293T cells (4.5 × 10^5^) were transfected with jetPRIME (Polyplustransfection, Illkirch, Strasbourg, France) following the manufacturer’s protocol. A total of 1 μg of each plasmid (eSpCas9(1.1) and HDR Vector) was used per well in a 6-well plate. For transfections that required more than one HDR vector, 700 ng of each plasmid was used. HCT116 or HELA cells (1 × 10^6^) were electroporated with a Neon electroporation system (Invitrogen, Carlsbad, California, USA) following the manufacturer’s recommendations. Seventy-two hours after transfection or electroporation, the cells were harvested and seeded on a 100 mm dish containing a selection antibiotic. The culture media was changed every 48 h, and the selection antibiotics were used at the following concentrations: puromycin at 3 μM, blasticidin at 10 μg/mL, and zeocin at 100 μg/mL.

### Validation of the integration of recombinant tags into the genomic DNA

Genomic DNA was extracted from modified cell lines using QuickExtract DNA extraction buffer (Epicentre Technologies, Madison, Wisconsin, USA) according to the manufacturer’s protocol. A set of genespecific primers were designed for each target using the NCBI primer design tool (https://www.ncbi.nlm.nih.gov/tools/primer-blast/) (Supplementary Fig. 1A and Supplementary table 2). The forward and reverse primers were designed to anneal approximately ^~^750 bp upstream and downstream, respectively, of the CRISPR targeting site of each gene under study. Additionally, a universal reverse primer that annealed to the linker of the HDR plasmid was included in every PCR mixture to generate a PCR product of ^~^750 bp if there was insertion of the recombinant DNA at the targeting site.

### Confocal live-cell microscopy

The Operetta CLS Confocal High Content Imaging System (PerkinElmer, Waltham, Massachusetts, USA) was used for single and timelapse image acquisition. The fluorescent proteins were detected with the following excitation and emission filter combinations: mTagBFP2 (Ex-405, Em-440), mClover3 (Ex-480, Em-513), and mRuby3 (Ex-558, Em-590). The images were acquired with a 20×, 40×, or 63× water immersion objective and 200–500 ms of acquisition time per fluorescent channel. All confocal images in this work were acquired with live cells grown in DMEM without phenol red (Thermo Fisher Scientific, Waltham, Massachusetts, USA) supplemented with 10% FBS in 96-or 384-well CellCarrier Ultra plates (PerkinElmer, Waltham, Massachusetts, USA). For time-lapse microscopy, cells were kept under environmentally controlled conditions (37°C, 5% CO_2_), and images were acquired every 10 min. All compounds under study were administered 50 min after the start of image acquisition at the following concentrations: staurosporine, 500 nM; paclitaxel, 20 nM; BI-6727, 1 μM; CX-6258, 1 μM.

### Airyscan superresolution microscopy

HEK293T cells with nuclear (H3.3-mRuby3), microtubule (β tubulin-mClover3), and mitochondrial (ATP5B-mTagBFP2) labeling were grown in phenol red-free DMEM supplemented with 10% FBS, 2 mM L-glutamine, 100 U/ml penicillin, and 100 μg/ml streptomycin at 37°C and 5% CO_2_. For Airyscan imaging, cells were plated onto No 1.5, 35 mm MatTek imaging chambers (MatTek, Ashland, Massachusetts, USA) pre-coated with 400-600 μg/ml Matrigel (Corning Corp, Corning, New York, USA), and cells were seeded to achieve ^~^60% confluency at the time of imaging. Cells were imaged in FluoroBrite DMEM (Thermo Fisher Scientific, Waltham, Massachusetts, USA) with 10% (v/v) FBS (Corning, Corning, New York, USA). Airyscan imaging was performed using a Zeiss 880 (Carl Zeiss AG, Oberkochen, Germany) outfitted with an Airyscan module and incubation chamber held at 37°C and 5% CO_2_. Data were collected using a 63× 1.4 NA objective and immersion oil optimized for 37°C (Carl Zeiss AG, Oberkochen, Germany). Colors were collected sequentially to minimize crosstalk and Airyscan processing was performed using the Airyscan module in the commercial ZEN software package (Carl Zeiss AG, Oberkochen, Germany). A rolling ball background subtraction with a radius of 50 pixels was subsequently applied in Fiji software (National Institute of Health, Bethesda, Maryland, USA).

### Luciferase live-cell detection

The NRF2-NanoLuc HCT116 cell line was used to evaluate the expression of NanoLuc-tagged NRF2 protein. For this purpose, 1 × 10^4^ cells in 100 μL of culture media were seeded into 96-or 384-well white plates, and 16 h later, the cells were treated with EndurenTM Live-cell luciferase substrate (Promega, Madison, Wisconsin, USA) at a final concentration of 40 μM. Ninety minutes later, the plates were transferred to a Tecan Spark plate reader (Tecan, Männerdorf, Switzerland) with environmental control (5% CO_2_, 37°C) to detect the luciferase signal. The luciferase signal of every well under study was tracked every 15 min with an integration time of 1 s per well, and the experiments were continued for 23 h. To evaluate the effect of terbutylhydroperoxide (TBHP, 25 μM), sulforaphane (SFN, 6.25 μM), and terbutylhydroquinone (TBHQ, 25 μM) on NRF2 expression, the cells were treated with the compounds 90 min after addition of the EndurenTM substrate and immediately transferred to the plate reader to start signal acquisition. Data were acquired with Spark control software and exported to Microsoft Excel or GraphPad Prism 6 for analysis.

### High-content imaging drug screening

The HELA cell line with multiplex endogenous labeling of the nucleus (Histone 1-mtagBFP2), microtubules (β tubulin-mClover3), and the autophagy receptor protein (SQSTM1-mRuby3) was used to perform high-content imaging drug screening of a small compound library composed of 352 kinase inhibitors (SelleckChem, Houston, Texas, USA). CellCarrier Ultra plates (384-well) were seeded with 5 × 10^3^ cells per well and 16 h later the cells were treated with each library compound at a final concentration of 1 μM. The effects of the compounds were evaluated 24 h later using an Operetta CLS Confocal High Content Imaging System (PerkinElmer, Waltham, Massachusetts, USA) with the following parameters: objective: 20 × water immersion; excitation and emission filter combination: mTagBFP2 (Ex-405, Em-440), mClover3 (Ex-480, Em-513), and mRuby3 (Ex-558, Em-590); acquisition time: 500 ms per channel. Two images (with three different channels) were acquired per well with the same coordinates in every well. As a control, 4 wells of cells received hydroxyl-chloroquine (25μM), a known inhibitor of autophagy and 28 wells of cells were treated only with DMSO (0.5% final concentration). The image analysis to determine cell nucleus localization, cytoplasm localization, and autophagic vesicles changes was performed with Harmony 4.7 software (PerkinElmer, Waltham, Massachusetts, USA).

### Reproducibility and Statistical Analysis

All figures in this work are representative of at least three independent experiments. Differences between control and treated samples were performed by using two-way analysis of variance in Graphpad Prism version 6.

## ACKNOWLEDGEMENTS

This work was supported in part by the National Institute of Health Grant R03DK105267.

## AUTHOR CONTRIBUTIONS

OPL conceived the project, designed the experiments, generated the vectors, transfected mammalian cells, analyzed the data, and wrote the paper. JNA performed super resolution microscopy experiments. JG performed high throughput compound screening experiments. CAB performed PCR experiments and MCR performed mammalian cell culture.

