## Supplementary-files for "A Versatile Vector System for the Fast Generation of Knock-in Cell Lines with CRISPR"

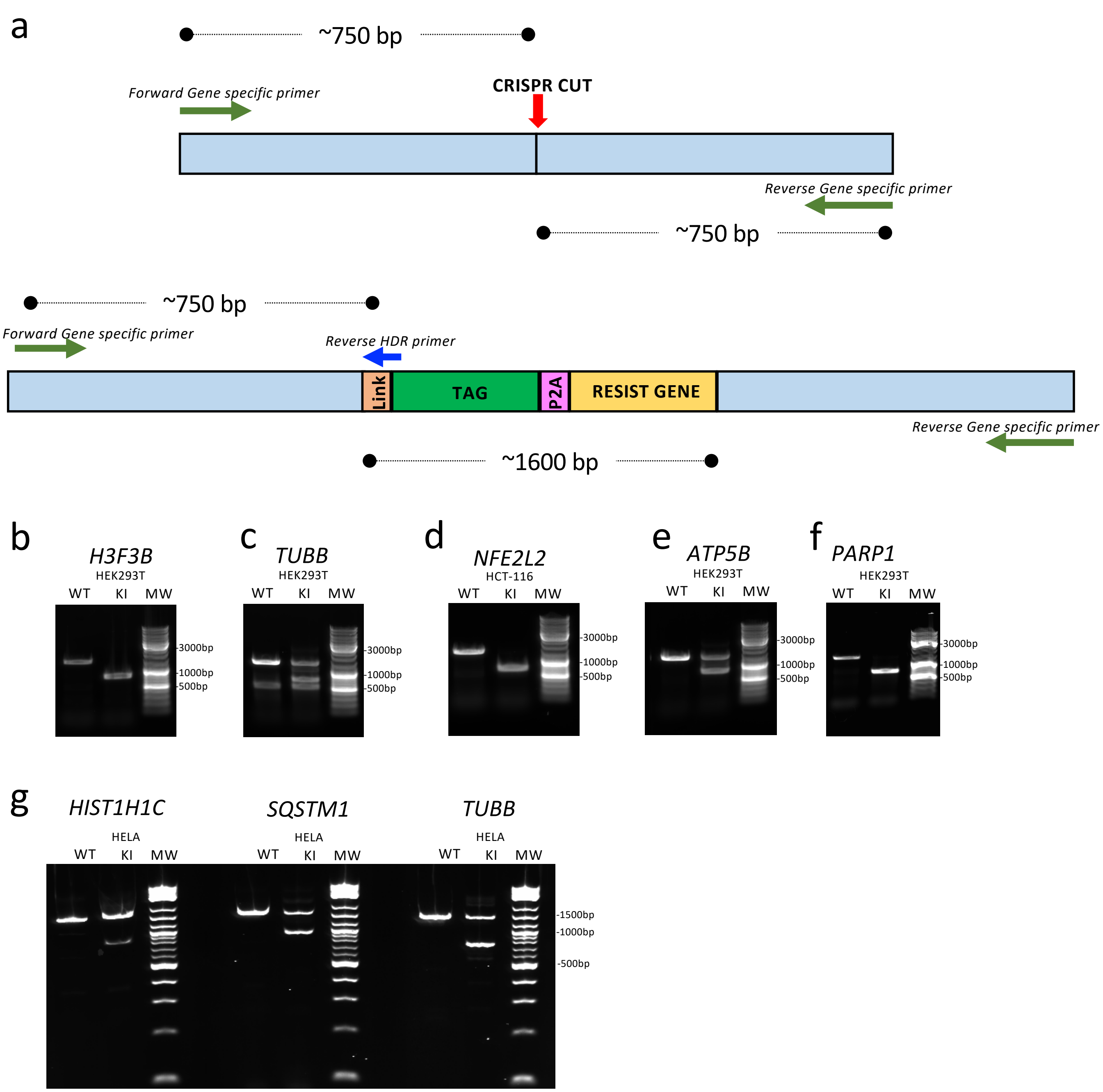


**Supplementary Figure 1**. Validation of genomic insertion of the labeling tags. (a) The genomic insertion was verified by PCR using a set of three primers. Two primers (green, forward and reverse) annealed around the desired CRISPR cutting site and generated a PCR product of ~1,500 bp if the region was not modified (upper panel). The third primer (reverse, in blue) produced a PCR product of ~750 bp only if the labeling tag was inserted at the desired location (lower panel). (b-g) The pattern of PCR products in wild-type and modified cells allowed us to determine the absence of modification or the presence of heterozygous or homozygous labeling. The correct targeting of the genes described in this work was validated as follows: (b) HEK293T cells with H3.3 labeled with mRuby3 (homozygous), (c) HEK293T cells with β tubulin labeled with mClover3 (Heterozygous), (d) HCT116 cells with NRF2 labeled with NanoLuc (homozygous), (e) HEK293T cells with ATP5B labeled with mTagBFP2 (heterozygous), (f) HEK293T cells with PARP1 labeled with mClover3 (homozygous), (g) HELA cells with multiplexing labeling of HIST1H1C with mTagBFP2 (Heterozygous), SQSTM1 with mRuby3 (Heterozygous) and TUBB with mClover3 (Heterozygous).

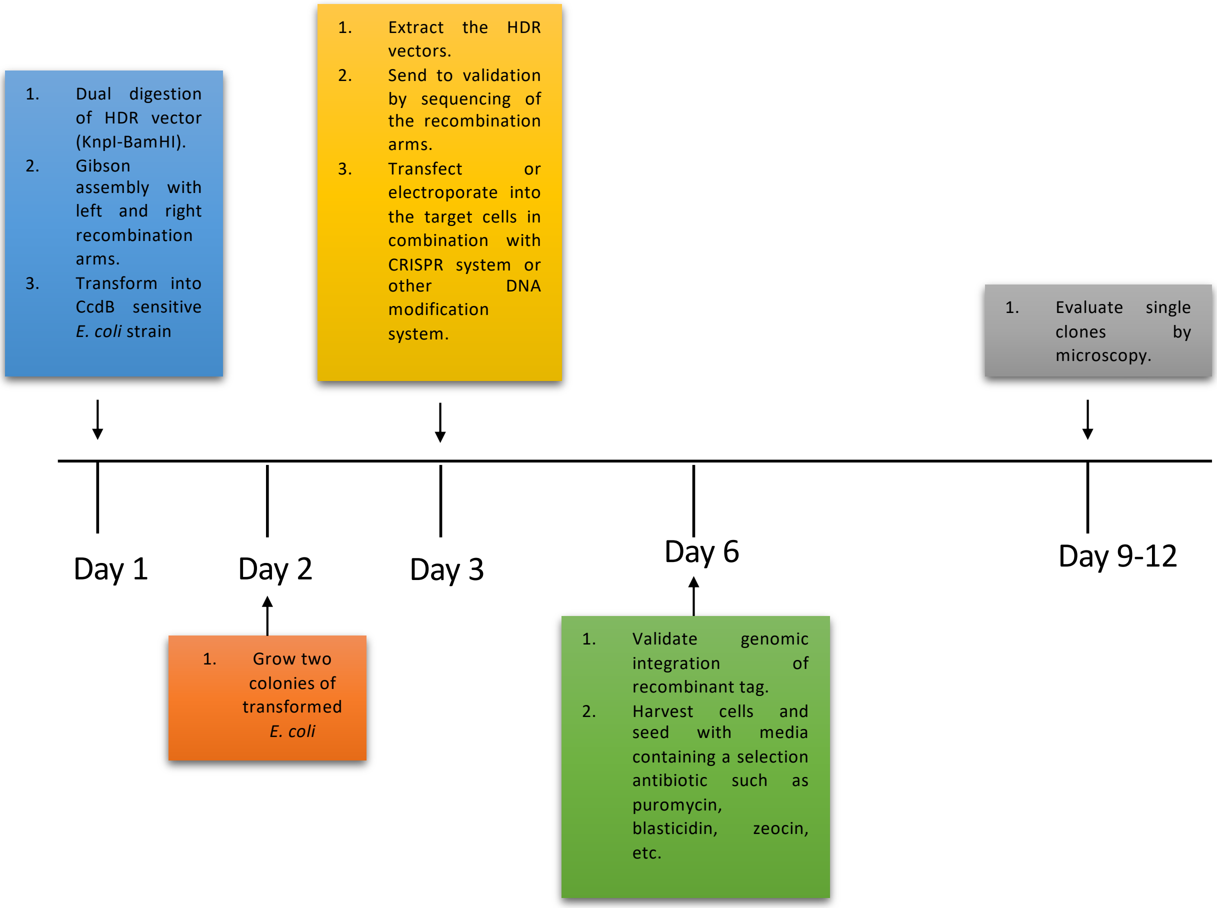


**Supplementary Figure 2**. Diagram of the process for developing homologous recombination vectors and selection of modified cell lines with the FAST-HDR system.

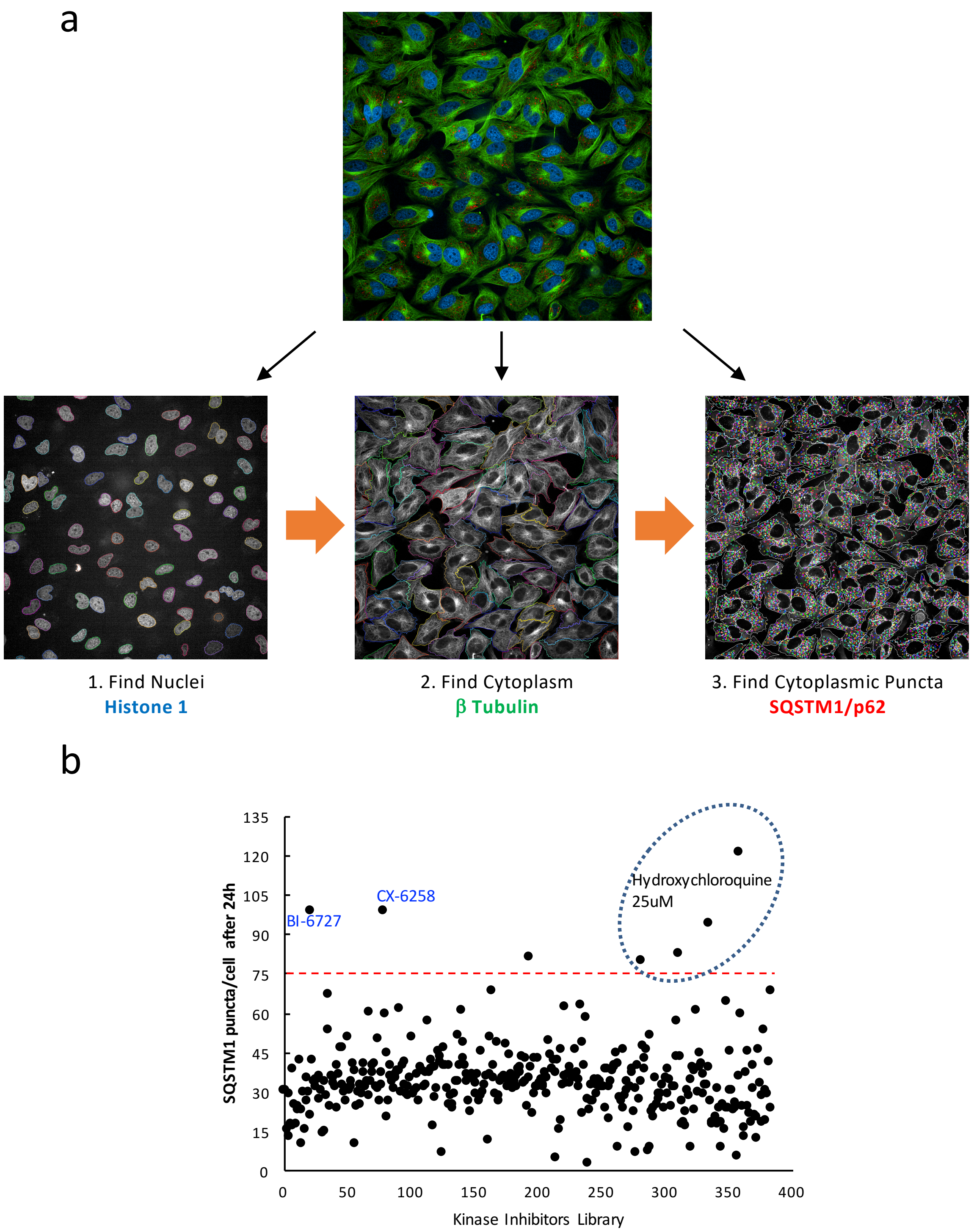


**Supplementary Figure 3.** Screening of a kinase inhibitor compound library for the identification of compounds that can induce the accumulation of cytoplasmic autophagic vesicles. (a) A HELA cell line with three endogenously tagged genes (Histone 1-mtagBFP2, Tubulin-Clover3 and p62/SQSTm1-mRuby3) was used to identify compounds that promote the accumulation of autophagic vesicles by using high-content imaging analysis. The analysis of the images of cells treated with every compound was performed with Harmony 4.7 software (Perkin Elmer) by following these steps: First, the location and quantitation of cells was done by finding the nuclear labeling of Histone 1-mtagBFP2, next the cytoplasm of each cell was identified by measuring the signal of Tubulin-mClover3, and lastly, the number of autophagic vesicles in the cytoplasm was determine by quantifying the number of puncta in the cytoplasm with the signal of p62/SQSTM1-mRuby3. (b) The average number of cytoplasmatic autophagic vesicles was quantitated in cells treated with 352 kinase inhibitors (1μM) in a 384 well-plate assay. Additionally, Hydroxychloroquine (25μM) or DMSO was used as positive and negative controls respectively. The red line defines the mean plus three standard deviation values. The top two compounds for inducing the accumulation of autophagic vesicles (BI-6727 and CX-6258) were selected for validation.

**Supplementary Table 1**. Sequences of the Synthetic DNA constructs for developing homologous recombination vectors for the genes targeted in this work. The Flanking sequences for Gibson assembly are colored in red.

| Gene Target | 5' Arm | 3' Arm | SgRNA sequence (underlined) |
|---|---|---|---|
| ATP5B (ATP synthase subunit B) | GACGTTGTAAAACGACGGCCA<br>GTGGGTACCCGCCACCACGCC<br>TGGCTAATTTTTGTATTTTTAGT<br>AGCCATGGGGTTTCACCATGTT<br>GGCTGGGCTTGTCTCGAACTC<br>CTGACCTTAGGTGATCCGCCT<br>GCCTTGGCCTCCCAAAGTGCT<br>GGGATTACAGGTGTAAGCCAC<br>CGTGCCTGGCCCATGTGTTCTT<br>AATTCATACTGTATCATATCTTG<br>TAAATTTGATTTGTGAGGGAAA<br>TTTAAGCTTTCTAAGATGACAT<br>GAATTCATCACATTCTAACTGA<br>TGGCCTGAAGTGGTGAGGAAT<br>GTTACATGATGCAGAAAGTTGA<br>TATCCCTCCGCTTCTTACTCTT<br>TTTTTTTTTCTCCCCCATCATAC<br>AGGTGAATATGACCATCTCCCA<br>GAACAGGCCTTCTATATGGTG<br>GGACCCATTGAAGAAGCTGTG<br>GCAAAAGCTGATAAGCTGGCT<br>GAAGAGCATTCATCGGGTACC<br>CAAGGCGGTGGAGAATTC | CAAGTCCCTGCGGTGTCTTTG<br>CTTGGATCCGAGGGGTCTTTG<br>TCCTCTGTACTGTCTCTCTCCT<br>TGCCCCTAACCCAAAAAGCTTC<br>ATTTTTCTGTGTAGGCTGCACA<br>AGAGCCTTGATTGAAGATATAT<br>TCTTTCTGAACAGTATTTAAGG<br>TTTCCAATAAAATGTACACCCC<br>TCAGAATTTGTCTGATTCTCTT<br>GGTTCTGACAACATAGTCAACA<br>CTGAAGGGTTATGTATTTAATT<br>TTAGTTTTAGAGACACGGTGTC<br>TGGCTGTGTTGCCAAGACTGG<br>TCTCTAACTCCTGGGCTCGAGA<br>TCTCCCACCTCAGTCTCCTGAG<br>TAGCTGGGGCTACAGGTGTAT<br>GTAGTCTCACATCACCAGCACT<br>GTTTTCAACAATTAGATTTTTAG<br>AGTGGCTATAAGAAGCAGTTTC<br>AGCATGAAGTGGGCCATGTAT<br>GTTTGAAATTGGTCCTTAAAAA<br>TAGCCATTCTCTGGATCCCGG<br>GCCCGTCGACTGCAGAGGCCT | GCGATCCGAGTTCAAATCTCG<br>GTGGAACCTTGAAGAGCATTCA<br>TCGTGAGGTTTTAGAGCTAGAA<br>ATAGCAAGTTAAAAT |
| HIST1H1C (Histone 1) | GACGTTGTAAAACGACGGCCA<br>GTGGGTACCCTGCCGCCGGCT<br>ATGATGTGGAGAAAAACAACAG<br>CCGTATCAAACTTGGTCTCAAG<br>AGCCTGGTGAGCAAGGGCACT<br>CTGGTGCAAACGAAAGGCACC<br>GGTGCTTCTGGCTCCTTTAAAC<br>TCAACAAGAAGGCAGCCTCCG<br>GGGAAGCCAAGCCCAAGGTTA<br>AAAAGGCGGGCGGAACCAAAC<br>CTAAGAAGCCAGTTGGGGCAG<br>CCAAGAAGCCCAAGAAGGCGG<br>CTGGCGGCGCAACTCCGAAGA<br>AGAGCGCTAAGAAAACACCGA<br>AGAAAGCGAAGAAGCCGGCCG<br>CGGCCACTGTAACCAAGAAAG<br>TGGCTAAGAGCCCAAAGAAGG<br>CCAAGGTTGCGAAGCCCAAGA<br>AAGCTGCCAAAAGTGCTGCTAA<br>GGCTGTGAAGCCCAAGGCCGC<br>TAAGCCCAAGGTTGTCAAGCCT<br>AAGAAA GCAGCA<br>CCTAAGAAGAAAGGTACCCAA<br>GGCGGTGGAGAATTC | CAAGTCCCTGCGGTGTCTTTG<br>CTTGGATCCAGGCGGCGCCCA<br>AGAAGAAATAGGCGAACGCCT<br>ACTTCTAAAACCCAAAAGGCTC<br>TTTTCAGAGCCACCACTGATCT<br>CAATAAAAGAGCTGGATAATTT<br>CTTTACTATCTGCCTTTTCTTGT<br>TCTGCCCTGTTACTTAAGGTTA<br>GTCGTATGGGAGTTACTGAGG<br>TATCAGACGAATTGGGTGACG<br>GGGTTGGAGAGTGGCCGTGGT<br>GAGGTTACAGCATTTAAACCTT<br>TATTGCGGCTTCTAGGTCCCTG<br>ACCGGAGGCTTTTCTCGCTGG<br>CGGATGGTTTTGGGATGGCAG<br>TCCCGCCCCAGGCCTGTGAAC<br>GGCAGAAAAGACCGCAAAACA<br>AGAGCCAGTTTCTTAGTCTAAA<br>GGGATGTCCGGATTGGACTAA<br>AAAATTTTCAAAAGTCCCGCCC<br>TGCTCCCGGGTTGGTCCGTTC<br>TTCTAGTACATGACTTTCAGGA<br>TCCCGGGCCCGTCGACTGCAG<br>AGGCCT | GCGATCCGAGTTCAAATCTCG<br>GTGGAACCTGGTTGTCAAGCC<br>TAAGAAGGGTTTTAGAGCTAGA<br>AATAGCAAGTTAAAAT |

| | | | |
|---|---|---|---|
| H3F3B (Histone 3.3) | GACGTTGTAAAACGACGGCCA<br>GTGGGTACCCGGCCTTATCTT<br>CGGGGCGTCTTTCTTAGGTGA<br>AAGAAAATGGCCCGAACCAAG<br>CAGACTGCTCGTAAGTCCACC<br>GGTGGGAAAGCCCCCCGCAAA<br>CAGCTGGCCACGAAAGCCGCC<br>AGGAAAAGCGCTCCCTCTACC<br>GGCGGGGTGAAGAAGCCTCAT<br>CGCTACAGGTAGGTCGGGCGG<br>GGGAACAATGGCCCGGCGGTG<br>GCCGGCTTTGTGCGGCAGCGT<br>CCGCTCACTCCTCCCCTGCTC<br>GCTGCAGGCCCGGGACCGTG<br>GCGCTTCGAGAGATTCGTCGT<br>TATCAGAAGTCGACCGAGCTG<br>CTCATCCGGAAGCTGCCCTTC<br>CAGAGGTTGGTGAGGGAGATC<br>GCGCAGGATTTCAAAACCGAC<br>CTGAGGTTTCAGAGCGCAGCC<br>ATCGGTGCGCTGCAGGTAAGA<br>CAAAGGCCTGGAGCCGGGGG<br>AGGGCTGGGCGGTTTCCGCTC<br>CCCGAGTGGGATTAATAGTGC<br>GGCTCTCGTCCTCAACAGGAG<br>GCTAGCGAAGCGTACCTGGTG<br>GGTCTGTTCGAAGATACCAACC<br>TGTGTGCCATCCACGCTAAGA<br>GAGTCACCATCATGCCCAAAG<br>ACATCCAGTTGGCTCGCCGGA<br>TAAGAGGAGAACGGGCGGGTA<br>CCCAAGGCGGTGGAGAATTC | CAAGTCCCTGCGGTGTCTTTG<br>CTTGGATCCACGGGGAGAGAG<br>AGCTTAAGTGAAGGCAGTTTTT<br>ATGGCGTTTTGTAGTAAATTCT<br>GTAAAATACTTTGGTTTAATTTG<br>TGACTTTTTTTGTAAGAAATTGT<br>TTATAATATGTTGCATTTGTACT<br>TAAGTCATTCCATCTTTCACTC<br>AGGATGAATGCGAAAAGTGAC<br>TGTTCACAGACCTCAGTGATGT<br>GAGCACTGTTGCTCAGGAGTG<br>ACAAGTTGCTAATATGCAGAAG<br>GGATGGGTGATACTTCTTGCTT<br>CTCATGATGCATGTTTCTGTAT<br>GTTAATGACTTGTTGGGTAGCT<br>ATTAAGGTACTAGAGTTGATAA<br>ATGTGTACAGGGTCCTTTTGCA<br>ATAAAACTGGTTATGACTTGAT<br>CCAAGTGTTTAACAATTGGGGC<br>TGTTAAGTCTGACCATACATCA<br>CTGTGATAGAATGTGGGCTTTT<br>TCAAGGGTGAAGATACAAGTCT<br>TAACCACAGTGTAACTTACAGT<br>TTCCTTTAAAAAAGGATCCCGG<br>GCCCGTCGACTGCAGAGGCCT | GCGATCCGAGTTCAAATCTCG<br>GTGGAACCTCAGTTGGCTCGC<br>CGGATACGGTTTTAGAGCTAGA<br>AATAGCAAGTTAAAAT |
| TUBB ( Tubulin) | GACGTTGTAAAACGACGGCCA<br>GTGGGTACCAGAACATGATGG<br>CTGCCTGTGACCCCCGCCACG<br>GCCGATACCTCACCGTGGCTG<br>CTGTCTTCCGTGGTCGGATGT<br>CCATGAAGGAGGTCGATGAGC<br>AGATGCTTAACGTGCAGAACAA<br>GAACAGCAGCTACTTTGTGGAA<br>TGGATCCCCAACAATGTCAAGA<br>CAGCCGTCTGTGACATCCCAC<br>CTCGTGGCCTCAAGATGGCAG<br>TCACCTTCATTGGCAATAGCAC<br>AGCCATCCAGGAGCTCTTCAA<br>GCGCATCTCGGAGCAGTTCAC<br>TGCCATGTTCCGCCGGAAGGC<br>CTTCCTCCACTGGTACACAGG<br>CGAGGGCATGGACGAGATGGA<br>GTTCACCGAGGCTGAGAGCAA<br>CATGAACGACCTCGTCTCTGA<br>GTATCAGCAGTACCAGGATGC<br>CACCGCAGAAGAGGAGGAGGA<br>TTTCGGTGAGGAGGCCGAAGA<br>GGAGGGTACCCAAGGCGGTG<br>GAGAATTC | CAAGTCCCTGCGGTGTCTTTG<br>CTTGGATCCCTAAGGCAGAGC<br>CCCCATCACCTCAGGCTTCTCA<br>GTTCCCTTAGCCGTCTTACTCA<br>ACTGCCCCTTTCCTCTCCCTCA<br>GAATTTGTGTTTGCTGCCTCTA<br>TCTTGTTTTTTGTTTTTTCTTCT<br>GGGGGGGGTCTAGAACAGTGC<br>CTGGCACATAGTAGGCGCTCA<br>ATAAATACTTGTTTGTTGAATGT<br>CTCCTCTCTCTTTCCACTCTGG<br>GAAACCTAGGTTTCTGCCATTC<br>TGGGTGACCCTGTATTTCTTTC<br>TGGTGCCCATTCCATTTGTCCA<br>GTTAATACTTCCTCTTAAAAATC<br>TCCAAGAAGCTGGGTCTCCAG<br>ATCCCATTTAGAACCAACCAGG<br>TGCTGAAAACACATGTAGATAA<br>TGGCCATCATCCTAAGCCCAAA<br>GTAGAAAATGGTAGAAGGTAGT<br>GGGTAGAAGTCACTATATAAGG<br>AAGGGGATGGGGATCCCGGG<br>CCCGTCGACTGCAGAGGCCT | GCGATCCGAGTTCAAATCTCG<br>GTGGAACCTGAGGCCGAAGAG<br>GAGGCCTAGTTTTAGAGCTAGA<br>AATAGCAAGTTAAAAT |

| | | | |
|---|---|---|---|
| NFE2L2 (Nuclear factor (erythroid-derived 2)-like 2 or NRF2) | GACGTTGTAAAACGACGGCCA<br>GTGGGTACCTTAGGGCAAAAG<br>CTCTCCATATCCCATTCCCTGT<br>AGAAAAAATCATTAACCTCCCT<br>GTTGTTGACTTCAACGAAATGA<br>TGTCCAAAGAGCAGTTCAATGA<br>AGCTCAACTTGCATTAATTCGG<br>GATATACGTAGGAGGGGTAAG<br>AATAAAGTGGCTGCTCAGAATT<br>GCAGAAAAAGAAAACTGGAAAA<br>TATAGTAGAACTAGAGCAAGAT<br>TTAGATCATTTGAAAGATGAAA<br>AAGAAAAATTGCTCAAAGAAAA<br>AGGAGAAAATGACAAAAGCCTT<br>CACCTACTGAAAAAACAACTCA<br>GCACCTTATATCTCGAAGTTTT<br>CAGCATGCTACGTGATGAAGAT<br>GGAAAACCTTATTCTCCTAGTG<br>AATACTCCCTGCAGCAAACAAG<br>AGATGGCAATGTTTTCCTTGTT<br>CCCAAAAGTAAGAAGCCAGAT<br>GTTAAGAAAAACGGTACCCAAG<br>GCGGTGGAGAATTC | CAAGTCCCTGCGGTGTCTTTG<br>CTTGGATCCAGGAGGATTTGA<br>CCTTTTCTGAGCTAGTTTTTTTG<br>TACTATTATACTAAAAGCTCCTA<br>CTGTGATGTGAAATGCTCATAC<br>TTTATAAGTAATTCTATGCAAAA<br>TCATAGCCAAAACTAGTATAGA<br>AAATAATACGAAACTTTAAAAA<br>GCATTGGAGTGTCAGTATGTTG<br>AATCAGTAGTTTCACTTTAACT<br>GTAAACAATTTCTTAGGACACC<br>ATTTGGGCTAGTTTCTGTGTAA<br>GTGTAAATACTACAAAAACTTA<br>TTTATACTGTTCTTATGTCATTT<br>GTTATATTCATAGATTTATATGA<br>TGATATGACATCTGGCTAAAAA<br>GAAATTATTGCAAAACTAACCA<br>CTATGTACTTTTTTTATAAATACT<br>GTATGGACAAAAAATGGCATTT<br>TTTATATTAAATTGTTTAGGATC<br>CCGGGCCCGTCGACTGCAGAG<br>GCCT | GCGATCCGAGTTCAAATCTCG<br>GTGGAACCTTAAGAAAAACTAG<br>ATTTAGGGTTTTAGAGCTAGAA<br>ATAGCAAGTTAAAAT |
| SQSTM1 (p62 protein) | GACGTTGTAAAACGACGGCCA<br>GTGGGTACCTATTTCAGTGTCC<br>ATTGATGGTTCTGCTTACACAC<br>CACCTGGCTGCCTGGTGTCGC<br>AGTGGCAGAGTTGAGCAGTGT<br>GAAAAAGACTGCTTGGCCCTTT<br>ACAGGGAAAGCAGGTCCACTG<br>TGGCCTGTGAGGACGAGAGCT<br>CTGGGCAGGCTCGGACACTGG<br>CAGACCCTGGTCCTGGCTGGC<br>CAAGGCAGCAGGGTATGTGTT<br>TCGGGTCACTCACAGGGCTCA<br>GCACCACTCCTCATGGCTTCCT<br>TACTGTTTCGGCAGAGGCTGA<br>CCCGCGGCTGATTGAGTCCCT<br>CTCCCAGATGCTGTCCATGGG<br>CTTCTCTGATGAAGGCGGCTG<br>GCTCACCAGGCTCCTGCAGAC<br>CAAGAACTATGACATCGGAGC<br>GGCTCTGGACACCATCCAATA<br>CAGCAAGCACCCCCCACCGCT<br>TGGTACCCAAGGCGGTGGAGA<br>ATTC | CAAGTCCCTGCGGTGTCTTTG<br>CTTGGATCCAGTATTCAAAGCA<br>TCCCCCGCCGTTGTGACCACT<br>TTTGCCCACCTCTTCTGCGTGC<br>CCCTCTTCTGTCTCATAGTTGT<br>GTTAAGCTTGCGTAGAATTGCA<br>GGTCTCTGTACGGGCCAGTTT<br>CTCTGCCTTCTTCCAGGATCAG<br>GGGTTAGGGTGCAAGAAGCCA<br>TTTAGGGCAGCAAAACAAGTGA<br>CATGAAGGGAGGGTCCCTGTG<br>TGTGTGTGTGCTGATGTTTCCT<br>GGGTGCCCTGGCTCCTTGCAG<br>CAGGGCTGGGCCTGCGAGACC<br>CAAGGCTCACTGCAGCGCGCT<br>CCTGACCCCTCCCTGCAGGGG<br>CTACGTTAGCAGCCCAGCACA<br>TAGCTTGCCTAATGGCTTTCAC<br>TTTCTCTTTTGTTTTAAATGACT<br>CATAGGTCCCTGACATTTAGTT<br>GATTATTTTCTGCTACGGATCC<br>CGGGCCCGTCGACTGCAGAGG<br>CCT | GCGATCCGAGTTCAAATCTCG<br>GTGGAACCTGGGATGCTTTGA<br>ATACTGGAGTTTTAGAGCTAGA<br>AATAGCAAGTTAAAAT |
| PARP1 (Poly [ADP-ribose] polymerase 1) | GACGTTGTAAAACGACGGCCA<br>GTGGGTACCGGCAGACAAGGA<br>TTAGAGGCTGTCCTGTAGTGTG<br>TCCCATGGTGACTGTTTCCCTT<br>CTGTGGTCCCTCCCTGTGCATA<br>GCCTGGCTTACAGGGTATGAG<br>CCTTCCCCCAGTTTCCAGAGG<br>ATGATCTCCTCTCCTCAGTCTG<br>CCTGAAGAAGACTTAGAGTAAC<br>TTTCAGGCTGGCATTGAGCATC<br>CTGCCAGCCCCGGGGAGATGA<br>GGCAACCCAGCCCCATGAAGA<br>GGCCTTAGAGTGACTTTCAGG<br>CTGGCACTGAGCGTCCTGCCA<br>GCCTGGGGGAGATGAGGCACA<br>TGTACATACCCTCTGTTGTATG<br>GCTGTTGGCTCCTTAACAAGCT<br>TCCCCTCAGGTACATTGTCTAT<br>GATATTGCTCAGGTAAATCTGA<br>AGTATCTGCTGAAACTGAAATT<br>CAATTTTAAGACCTCCTTATGG<br>GGTACCCAAGGCGGTGGAGAA<br>TTC | CAAGTCCCTGCGGTGTCTTTG<br>CTTGGATCCCTGTGGTAATTGG<br>GAGAGGTAGCCGAGTCACACC<br>CGGTGGCTCTGGTATGAATTCA<br>CCCGAAGCGCTTCTGCACCAA<br>CTCACCTGGCCGCTAAGTTGC<br>TGATGGGTAGTACCTGTACTAA<br>ACCACCTCAGAAAGGATTTTAC<br>AGAAACGTGTTAAAGGTTTTCT<br>CTAACTTCTCAAGTCCCTTGTT<br>TTGTGTTGTGTCTGTGGGGAG<br>GGGTTGTTTTGGGGTTGTTTTT<br>GTTTTTTCTTGCCAGGTAGATA<br>AAACTGACATAGAGAAAAGGCT<br>GGAGAGAGATTCTGTTGCATA<br>GACTAGTCCTATGGAAAAAACC<br>AAGCTTCGTTAGAATGTCTGCC<br>TTACTGGTTTCCCCAGGGAAG<br>GAAAAATACACTTCCACCCTTT<br>TTTCTAAGTGTTCGTCTTTAGTT<br>TTGATTTTGGAAAGGATCCCGG<br>GCCCGTCGACTGCAGAGGCCT | GCGATCCGAGTTCAAATCTCG<br>GTGGAACCTTCAATTTTAAGAC<br>CTCCCTGGTTTTAGAGCTAGAA<br>ATAGCAAGTTAAAAT |

**Supplementary Table 2**. Sequences of primers used in this work.

| Genes | | Sequences (5'-3') |
|---|---|---|
| *ATP5B* | Forward | TGCTGTGGTCCCATTCCAACA |
| | Reverse | ACGAGTTTTTCCCAATATGCCCTTC |
| *HIST1H1C* | Forward | CCAGAGCACCAATCAGAGCG |
| | Reverse | CCTGTCAGTTTAGCGGAAGGC |
| *H3F3B* | Forward | CGCAGCCTGAGTCATTAGGGG |
| | Reverse | TGCAAGGTATAAATGCGCATAGCA |
| *TUBB* | Forward | GCACTCTGAAGCTGACCACACCA |
| | Reverse | CCTCCCAACCCCCTTGATCCCTT |
| *NFE2L2* | Forward | GTGCCCCTGGAAGTGTCAAACA |
| | Reverse | GTGGGCGTATGTCTACTGATGGAA |
| *SQSTM1* | Forward | TCTTTGCATCATCATAGCTTAGCATCT |
| | Reverse | GTCATTGGTTAAAGTGCTGATGCC |
| *PARP1* | Forward | CCCTGGAAGCTTTGCCACATC |
| | Reverse | AGCCCTTGGGTAAGTATATTTGTGG |
| *Universal Reverse Link* | Reverse | ATGAATTCTCCACCGCCTTG |
